# CREBBP/EP300 acetyltransferase inhibition disrupts FOXA1-bound enhancers to inhibit the proliferation of ER+ breast cancer cells

**DOI:** 10.1101/2021.12.26.474204

**Authors:** Archana Bommi-Reddy, Sungmi Park-Chouinard, David N. Mayhew, Esteban Terzo, Aparna Hingway, Michael J. Steinbaugh, Jonathan E. Wilson, Robert J. Sims, Andrew R. Conery

## Abstract

Therapeutic targeting of the estrogen receptor (ER) is a clinically validated approach for estrogen receptor positive breast cancer (ER+ BC), but sustained response is limited by acquired resistance. Targeting the transcriptional coactivators required for estrogen receptor activity represents an alternative approach that is not subject to the same limitations as targeting estrogen receptor itself. In this report we demonstrate that the acetyltransferase activity of coactivator paralogs CREBBP/EP300 represents a promising therapeutic target in ER+ BC. Using the potent and selective inhibitor CPI-1612, we show that CREBBP/EP300 acetyltransferase inhibition potently suppresses in vitro and in vivo growth of breast cancer cell line models and acts in a manner orthogonal to directly targeting ER. CREBBP/EP300 acetyltransferase inhibition suppresses ER-dependent transcription by targeting lineage-specific enhancers defined by the pioneer transcription factor FOXA1. These results validate CREBBP/EP300 acetyltransferase activity as a viable target for clinical development in ER+ breast cancer.

## INTRODUCTION

Estrogen Receptor positive breast cancer (ER+ BC) constitutes approximately 60-80% of breast cancer cases and ER signaling is acknowledged as the oncogenic driver of the disease (Rosenberg, Barker et al. 2015). As such, anti-estrogen therapy is the mainstay of treatment in ER+ BC (Ali, Buluwela et al. 2011). Although there is clear evidence that current therapies have prolonged patient survival, a majority of patients with metastatic disease acquire resistance and relapse (Osborne and Schiff 2011, Caswell-Jin, Plevritis et al. 2018). Therefore, new therapies are continually required to improve clinical outcomes for these patients.

Cyclic AMP response element-binding binding protein (CREBBP) and its closely related homolog E1A binding protein of 300 kDa (EP300) are ubiquitously expressed multidomain transcriptional coactivators whose histone acetyltransferase (HAT) domains are highly conserved throughout evolution (Wang, Tang et al. 2008). CREBBP/EP300 catalyze lysine acetylation on a broad range of substrates, particularly K18 and K27 on histone H3 and several non-histone proteins, to regulate signaling pathways involved in cell growth, development and tumorigenesis (Dancy and Cole 2015, Weinert, Narita et al. 2018). Recently, it has been demonstrated that CREBBP/EP300 act to regulate lineage-specific transcriptional programs (as opposed to global transcriptional programs), which are largely driven by distal enhancers and tissue-specific transcription factors (Narita, Ito et al. 2021).

CREBBP/EP300 are core components of the ER transcriptional complex (Yi, Wang et al. 2015), and acetyltransferase activity of CREBBP/EP300 is critical for ER signaling (Yi, Wang et al. 2017). Within the complex, CREBBP/EP300 act through direct acetylation of ER to enhance its DNA binding and transactivation function (Kim, Woo et al. 2006). Beyond this direct activity on ER, one might also hypothesize that ER-bound CREBBP/EP300 act on histones to create regions of hyperacetylation and open chromatin to facilitate the recruitment of transcriptional machinery. ER signaling is known to be reprogrammed through enhancer remodeling during breast tumorigenesis (Chi, Singhal et al. 2019). Further, the majority of ER binding sites (estrogen response elements, or ERE) have been mapped to distal enhancers, consistent with the activity of CREBBP/EP300 (Lin, Vega et al. 2007, Welboren, van Driel et al. 2009). The magnitude of changes in chromatin accessibility and the relationship to the histone acetyltransferase activity of CREBBP/EP300 have not been examined in the context of ER-driven transcription.

We propose that inhibiting the HAT domain of CREBBP/EP300 will be an effective strategy to orthogonally target the clinically validated ER transcriptional network. By not targeting ER directly, therapeutic inhibition of CREBBP/EP300 has the advantage of being active in the context of resistance to anti-estrogen therapies. For many years the only options for targeting CREBBP/EP300 acetyltransferase activity were natural products or nonspecific inhibitors, but recently multiple highly selective, potent, and orally bioavailable CREBBP/EP300 HAT inhibitors have been described, including A-485 (Lasko, Jakob et al. 2017) and CPI-1612 (Wilson, Patel et al. 2020). Here we use CPI-1612 to demonstrate that inhibition of CREBBP/EP300 HAT activity represents a promising strategy to suppress ER-mediated proliferation and tumor growth. Further, we use this potent and selective inhibitor to provide insight into the role of CREBBP/EP300 in defining the transcriptional programs and chromatin landscape of ER+ breast cancer.

## RESULTS

### CREBBP/EP300 HAT inhibition abrogates ER-driven proliferation *in vitro* and *in vivo*

Given the functional link between CREBBP/EP300 and ER, we investigated whether inhibition of CREBBP/EP300 acetyltransferase activity would impact the viability of ER+ breast cancer cell lines. Pooled, barcoded breast cancer cell lines were treated with a dose titration of the previously described potent and selective inhibitor of CREBBP/EP300 acetyltransferase activity, CPI-1612 (Wilson, Patel et al. 2020). Growth inhibition was measured using barcode depletion after 5-day treatment with compound as described (Yu, Mannan et al. 2016). Consistent with previous results (Lasko, Jakob et al. 2017), CREBBP/EP300 HAT inhibition shows activity across multiple breast cancer cell lines, but notably the most sensitive cell lines were ER positive (Fig S1A).

We next confirmed that CPI-1612 is highly active in ER positive breast cancer cell lines in standard growth inhibition assays, with GI_50_ values below 100 nM (Fig 1A). Importantly, these GI_50_ values are in the same range as published EC_50_ values for reduced H3K18 acetylation by CPI-1612, arguing for on-target effects of CREBBP/EP300 inhibition (Wilson, Patel et al. 2020). To exclude impacts of CPI-1612 on non-hormone driven growth factor networks in full serum, we made use of growth factor/hormone depleted charcoal-stripped media and exogenously added estradiol (CSS+E2). We noted that GI_50_ values for CPI-1612 were similar for ER positive breast cancer cell lines in both culture conditions (Fig S1B), demonstrating that CREBBP/EP300 inhibition can directly impact hormone-driven proliferation.

**Figure 1.**
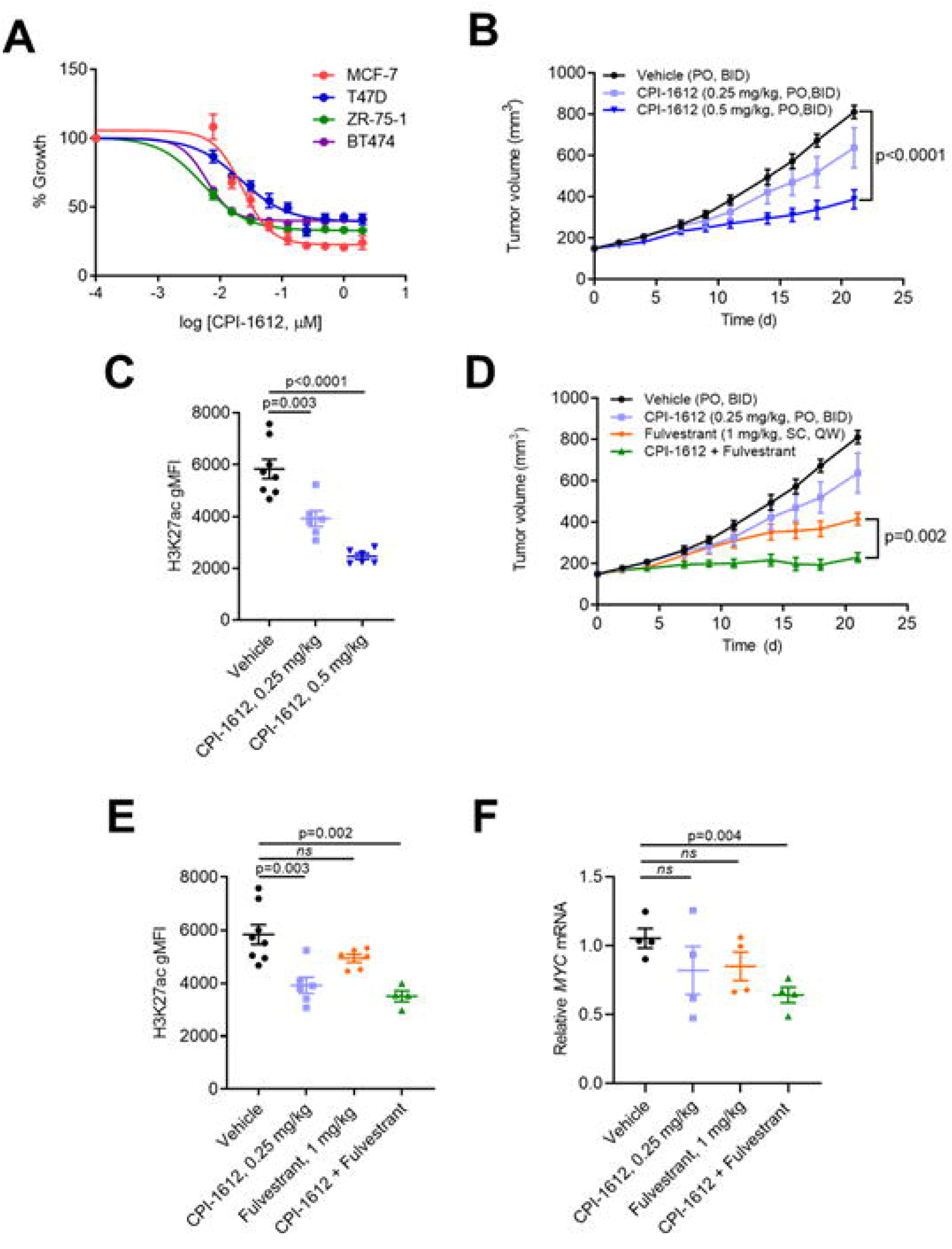
CPI-1612 inhibits viability of ER+ breast cancer cell lines and ER signaling both in vitro and in vivo. **A,** In vitro activity of CPI-1612. ER+ breast cancer cell lines were treated with increasing doses of CPI-1612 and cell viability was measured using Cell Titer Glo after 4 days of treatment. Error bars represent standard deviation (n=2). **B,** In vivo activity of single agent CPI-1612. Female Balb/c nude mice were implanted subcutaneously with MCF7 cells (n=8 for vehicle, n=6 for others) and treated with the indicated doses of CPI-1612 (PO, BID) or an equal volume of vehicle (PO, BID). Tumor volumes were measured by caliper until study termination at 21 days. Data points represent mean and SEM at each timepoint. P-values were calculated using student’s t-test relative to the vehicle arm; the p-value at study endpoint is shown (no data point in the 0.25 mg/kg arm reached statistical significance). **C,** Pharmacodynamic readout of CPI-1612 activity. PBMCs were isolated from blood at study termination, fixed, and stained for FACS analysis. The level of H3K27ac was quantified using gMFI (geometric mean fluorescence intensity). Data are represented as mean ± SEM, and p-values were calculated using student’s t-test. **D,** Efficacy of CPI-1612 in combination with Fulvestrant. Mice were xenografted with MCF7 cells as in **B**, and treated with CPI-1612 (0.25 mg/kg, PO, BID), Fulvestrant (1 mg/kg, SC, QW), CPI-1612 + Fulvestrant, or vehicle (n=8 for vehicle, n=6 for others). Data points represent the mean and SEM of surviving animals, and p-values were calculated at each timepoint using student’s t-test. P-value for the difference between Fulvestrant and CPI-1612 + Fulvestrant at study endpoint is shown. **E,** PD in PBMCs for study described in **D**, as in **C**. **F,** Tumor PD as measured by gene expression changes. Total mRNA was isolated from tumors collected at study endpoint and used for q-RTPCR analysis. *MYC* expression normalized to *ACTB* was calculated relative to vehicle mean and is expressed as mean ± SEM for each arm. P-values were calculated by student’s t-test relative to vehicle.

To investigate the anti-tumor effects of CREBBP/EP300 HAT inhibition in ER+ breast cancer in vivo, we used an MCF7 xenograft model. CPI-1612 has favorable ADME properties and achieves sufficient exposure to induce pharmacodynamic and anti-tumor responses in vivo (Wilson, Patel et al. 2020). Oral dosing with CPI-1612 either twice daily in established MCF7 xenografts resulted in dose-dependent inhibition of tumor growth (Fig 1B), and all doses were well tolerated (Figs S1C and S1D). Notably, CPI-1612 treatment led to dose-dependent reduction in H3K27 acetylation in peripheral blood, demonstrating target engagement at efficacious doses (Fig 1C).

Standard of care (SOC) treatment for ER positive breast cancers consists of anti-hormone therapy such as the selective ER degrader (SERD) Fulvestrant. To determine whether CREBBP/EP300 acetyltransferase inhibition could enhance the response to SOC therapy, we treated xenografted MCF7 cells with Fulvestrant and CPI-1612 alone or in combination. To maximize the potential for combinatorial effects, a suboptimal dose of CPI-1612 was used in combination with a clinically relevant dose of Fulvestrant (Bihani, Patel et al. 2017). As shown in Fig 1D, treatment with low dose CPI-1612 enhanced the anti-tumor efficacy of Fulvestrant in a well-tolerated dosing regimen that was associated with reduction in H3K27 acetylation (Fig 1E and S1E). Enhanced anti-tumor efficacy was also associated with a more pronounced reduction in the mRNA level of the known ER target MYC in tumor tissue, suggesting enhanced engagement of the ER transcriptional network (Fig 1F). As shown in Figs S1F and S1G, enhanced efficacy and pharmacodynamics were not the result of increased exposure of either CPI-1612 or Fulvestrant. Taken together, these data demonstrate that selective inhibition of CREBBP/EP300 acetyltransferase activity blocks the proliferation of ER positive breast cancer cells in vitro and in vivo, and the enhanced efficacy in combination with Fulvestrant suggests that acetyltransferase inhibition has the potential to potentiate the effects of direct ER targeting, perhaps through increased engagement of ER transcriptional programs.

### CREBBP/EP300 HAT inhibition inhibits ER-dependent transcriptional programs

To explore the transcriptional events underlying the phenotypic response to CREBBP/EP300 acetyltransferase inhibition, we carried out RNA sequencing analysis of three breast cancer cell lines (MCF7, T47D, and ZR751) treated with CPI-1612 alone or in combination with Fulvestrant. To minimize potential secondary effects of long-term treatment, we treated cells for 6 hours prior to sample preparation. As shown in Fig 2A and Fig S2A, CPI-1612 treatment has broad, dose-dependent effects on gene expression, with the majority of differentially expressed genes showing downregulation, while Fulvestrant has a modest transcriptional impact. Gene set enrichment analysis (GSEA) with Hallmark gene sets revealed that the top two downregulated gene signatures for both CPI-1612 and Fulvestrant treatment were HALLMARK_ESTROGEN_RESPONSE_EARLY and HALLMARK_ESTROGEN _RESPONSE_LATE, but CPI-1612 has a broader impact, consistent with the higher number of genes modulated (Figs 2B and S2B). Comparison of the enrichment plots for single agent CPI-1612 or Fulvestrant with combination treatment shows enhanced repression upon combination treatment (Fig S2C). Investigation of the genes driving gene set repression demonstrates that while both CPI-1612 and Fulvestrant target the ER transcriptional network, they do so by targeting non-identical sets of genes (Fig 2C). For example, while known ER target genes such as *MYC* and *GREB1* are regulated by both treatments, CPI-1612 uniquely regulates a set of genes including *KLF4, ELF3*, and *HES1* (Figs 2C, 2D, and S3A). As shown in Fig S3B, while genes that are acutely downregulated by Fulvestrant are largely defined by the presence of a proximal ERE, genes acutely downregulated by CREBBP/EP300 HAT inhibition appear to be defined by alternative features.

**Figure 2.**
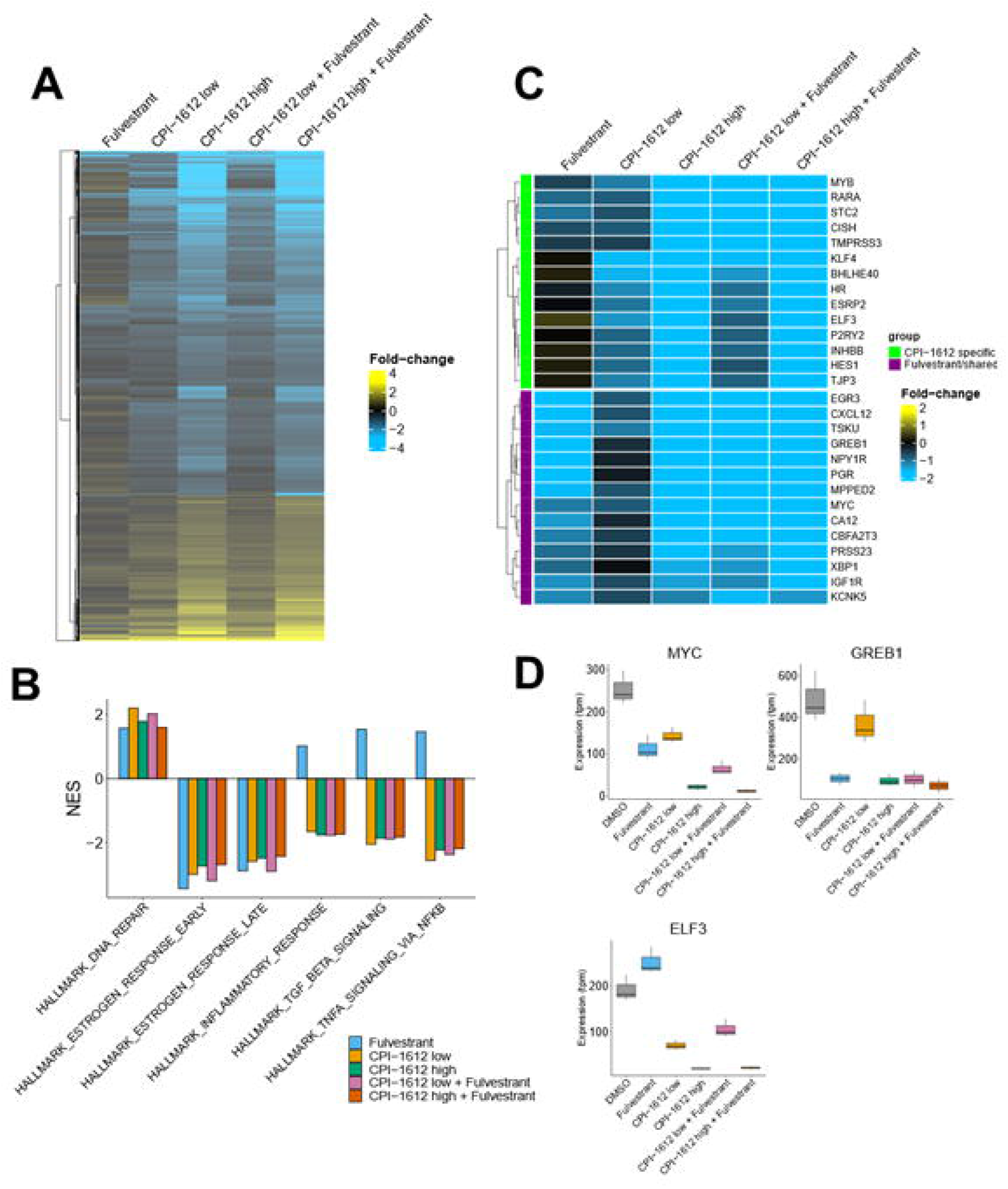
CPI-1612 inhibits the ER transcriptional program. **A,** Bulk RNA-seq analysis of MCF7 cells. MCF7 cells were treated as indicated for 6 hours (n=3 per treatment) followed by isolation of mRNA for RNA-sequencing analysis. Differential expression is indicated as log_2_ (fold-change) in normalized counts relative to the DMSO control. Concentrations used were CPI-1612 low: 5 nM; CPI-1612 high: 50 nM; Fulvestrant: 100 nM. **B,** CPI-1612 has a distinct transcriptional effect from Fulvestrant. Gene Set Enrichment Analysis (GSEA) was carried out using the MSigDB Hallmark genesets with the data described in **A**. Normalized enrichment scores (NES) for selected genesets are shown. **C,** CPI-1612 regulates the ER transcriptional network by impacting different genes than Fulvestrant. Subset of data in **A** showing selected genes in the HALLMARK_ESTROGEN_RESPONSE_EARLY geneset. Genes that are regulated by CPI-1612, Fulvestrant, or both are highlighted. **D,** Example of differentially regulated genes. Expression values from RNA-seq are quantified as transcripts per million (tpm) and are plotted from the experiment described in **A**. Boxplots show the median and range of tpm values from triplicate samples for each treatment.

### CREBBP/EP300 inhibition targets a subset of enhancers that are linked to differentially expressed genes

To better understand how CREBBP/EP300 HAT inhibition leads to transcriptional changes in ER+ breast cancer cells, we performed ATAC-seq and H3K27 acetyl ChIP-seq upon treatment of MCF7 cells with CPI-1612. As expected from published substrate profiling of CREBBP/EP300 (Jin, Yu et al. 2011, Weinert, Narita et al. 2018) and from the data shown in Fig 1C, CPI-1612 led to a profound loss of H3K27ac, with more than one-third of all peaks showing at least a two-fold decrease in CPI-1612 treated cells relative to the DMSO control (Figs 3A and 3B). As a control, there were minimal effects on H3K9ac, arguing for the selectivity of CPI-1612. Intriguingly, ATAC-seq results showed that effects on chromatin accessibility were much more modest than changes in H3K27ac, with only about 2% of all open peaks showing the same two-fold change in magnitude, with most differential peaks showing a reduction in accessibility (Figs 3A and 3B). As expected, total ATAC-seq peaks show a high overlap with total H3K27ac peaks (Fig S4A). However, while the majority of differential ATAC-seq peaks also show differential H3K27ac, the converse is not true, as <5% of differential H3K27ac peaks show significant changes in chromatin accessibility (Figs S4C and S4D). Thus, while CREBBP/EP300 HAT inhibition globally reduces H3K27 acetylation in ER+ breast cancer cells, changes in chromatin accessibility are much more circumscribed.

**Figure 3.**
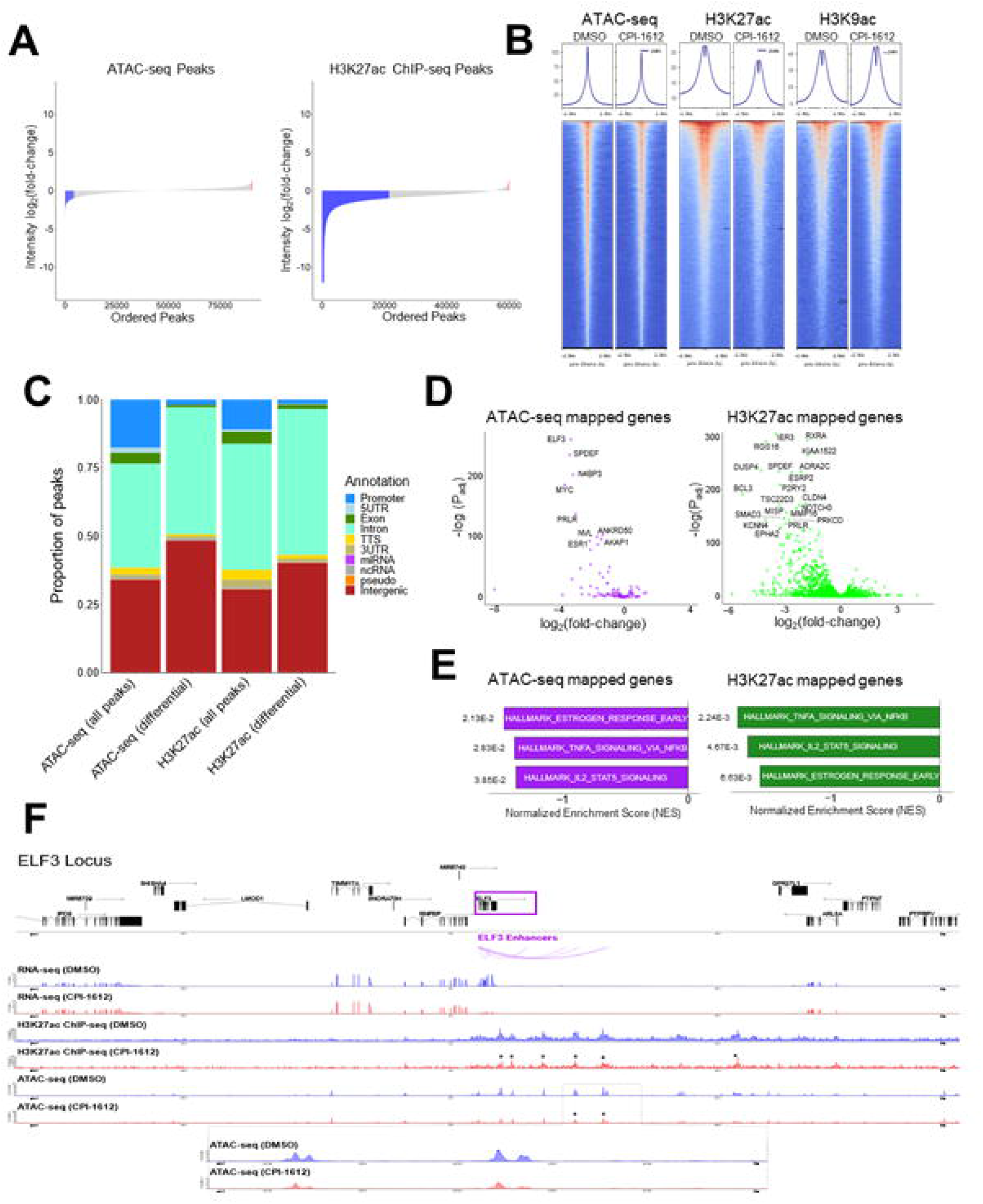
CPI-1612 represses a subset of enhancers which are linked to differentially expressed genes. **A,** CPI-1612 treatment impacts chromatin accessibility and histone acetylation. MCF7 cells were treated for 6 hours with DMSO or CPI-1612 (50 nM), and samples were prepared for ATAC-seq or ChIP-seq with H3K27ac or H3K9ac antibodies. Waterfall plot of the log_2_ (foldchange) in signal intensity in open chromatin (ATAC-seq) peaks and H3K27ac peaks between DMSO and 50 nM CPI-1612 treatment. Blue: peaks with at least 2-fold decrease in signal; red: peaks with at least 2-fold increase in signal. **B,** Global effects of CPI-1612 treatment. Top 30,000 peaks for each feature were ranked based on intensity for DMSO and CPI-1612 conditions. Graphs represent the sum of signal intensity across all peaks. C, Fraction of peaks located in different genomic regions. Peaks from ATAC-seq or H3K27ac were assigned to the indicated genomic regions using HOMER. All: all peaks identified in DMSO and CPI-1612 conditions; differential: peaks that changed at least 2-fold upon treatment. D, Genes mapped to differential ATAC-seq or H3K27ac peaks are likely to be downregulated. Genes were assigned to differential ATAC-seq or H3K27ac peaks, and differential expression data (as described in Fig 2) from DESeq2 were plotted. E, Genes linked to differential features are enriched for ER targets. Genes from D were used for GSEA with Hallmark genesets. Top three signatures are shown with NES on the x-axis and adjusted P-value (Padj) next to bars. F, Integrated gene expression and epigenomic features for the ELF3 locus. Top panel shows annotated genes, and purple lines show annotated ELF3 enhancer elements (Andersson, Gebhard et al. 2014). RNA-seq, H3K27ac ChIP-seq, and ATAC-seq tracks are plotted at the bottom, with H3K27ac and ATAC-seq peaks showing at least a 2-fold decrease marked with a *. The inset shows the differential ATAC-seq peaks in the ELF3 enhancer.

To determine if differentially accessible and acetylated sites were enriched in specific genomic regions, we annotated peaks with nearby genomic features using HOMER (Heinz, Benner et al. 2010). Both H3K27ac and ATAC-seq differential peaks were more likely to be found in intergenic regions and less likely to be found in promoters (Fig 3C). We next checked whether the genes linked to these enhancers were differentially expressed after CPI-1612 treatment by mapping peaks to genes using an approach described previously (Corces, Granja et al. 2018). Given the relatively smaller number of peaks, the ATAC-seq mapped gene list was smaller than the H3K27ac ChIP-seq mapped list (Fig 3D), yet both gene sets identify several key genes which were down-regulated after CPI-1612 treatment, including *MYC* and *ESR1*. GSEA of the genes mapped to differential ATAC-seq or H3K27ac peaks further showed enrichment of the ER transcriptional network (Fig 3E). The transcription factor *ELF3*, a member of the core transcriptional regulatory circuitry in MCF7 cells (Saint-Andre, Federation et al. 2016) was one of the most robustly down-regulated genes and is illustrative of the broad reduction of H3K27ac and more focal reduction in chromatin accessibility at distal enhancer sites (Fig 3F).

### CREBBP/EP300 inhibition targets FOXA1 cell type-specific binding sites that control ER signaling and luminal-specific gene sets in breast cancer cells

The large-scale decrease in H3K27ac signal was in line with expectations for inhibition of CREBBP/EP300, but it was less intuitive as to why a much smaller subset of enhancers would change chromatin accessibility status when profiled by ATAC-seq. To test whether these regions contained the motifs for any specific transcription factors, we employed HOMER’s findMotifs function (Heinz, Benner et al. 2010) to test whether the underlying DNA sequences of the subset of ATAC-seq peaks which close upon CPI-1612 treatment were enriched for any known transcription factor motifs. Strikingly, the FOXA1 transcription factor motif was the most statistically significant motif in this region, with other FOX family motifs also showing significant enrichment, likely due to sequence similarity (Fig 4A). In contrast, differential H3K27ac peaks did not show enrichment of FOXA1 motifs (Fig S5A). FOXA1 is a pioneer transcription factor that can open chromatin and facilitate the binding of other transcription factors, including ER, in breast cancer cell lines (Carroll, Liu et al. 2005, Hurtado, Holmes et al. 2011). To further confirm these observations, we independently performed single-cell ATAC-seq in MCF7 cells treated with either CPI-1612 or DMSO. The single-cell analysis revealed some heterogeneity in the epigenetic composition of MCF7 cells (Fig S6A); however, in concordance with the bulk ATAC-seq data, most regions of open chromatin did not change after treatment (Fig S6B). We used chromVAR (Schep, Wu et al. 2017) to identify the transcription factor motifs that had the largest change within variable peaks after treatment. Commensurate with the bulk ATAC-seq data, the motif for FOXA1 showed the largest decrease in predicted binding activity after treatment (Figs S6C and S6D).

**Figure 4.**
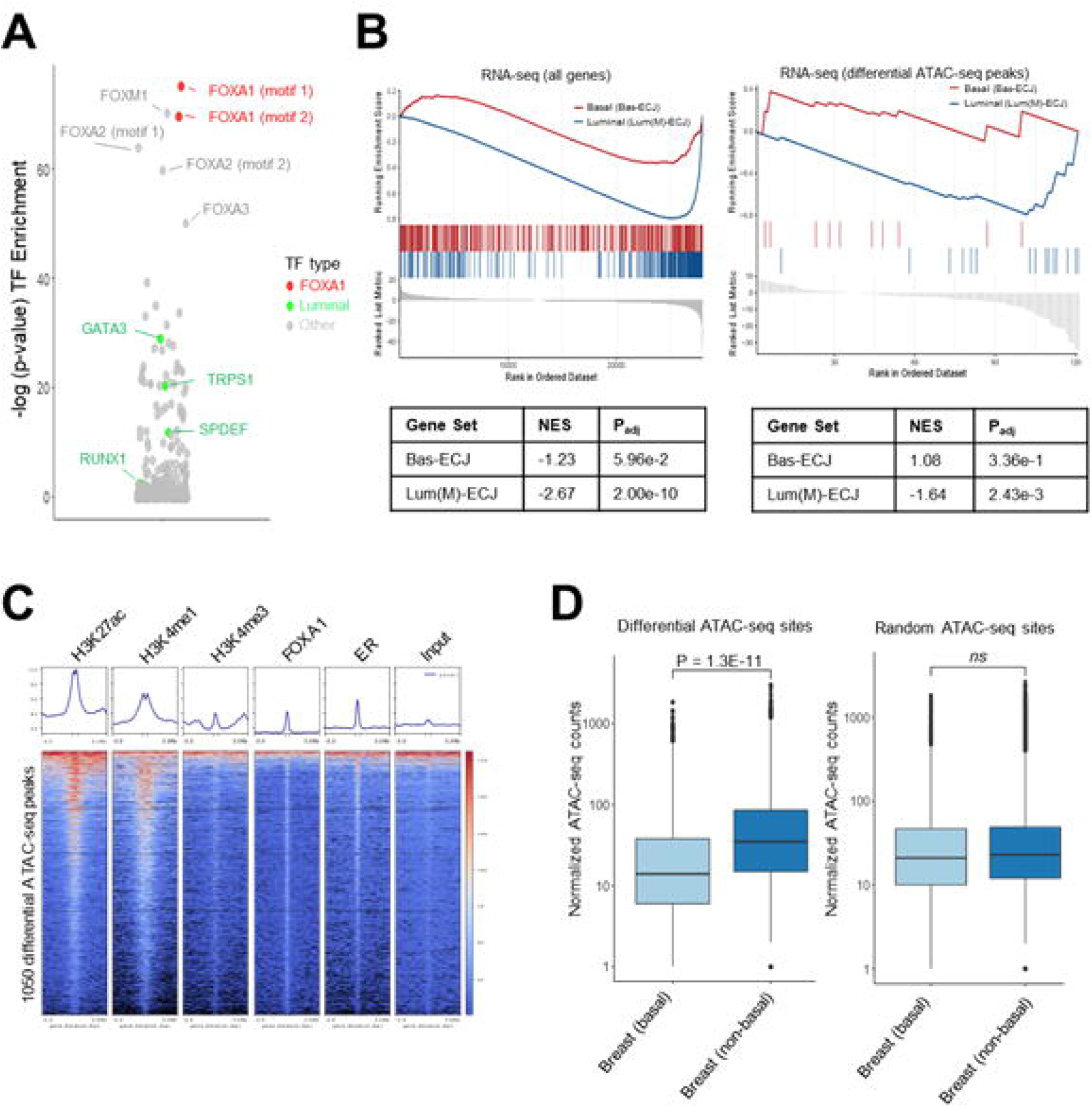
CPI-1612 targets FOXA1 binding sites that control luminal-specific gene sets in MCF7 cells and breast tumors. **A,** Differential ATAC-seq peaks are enriched for FOXA1 motifs. HOMER motif analysis was used to identify enrichment of transcription factor (TF) motifs in the ATAC-seq peaks that were downregulated at least 2-fold after CPI-1612 treatment, relative to the fraction of all ATAC-seq peaks with binding sites. **B,** Downregulated genes and genes mapped to sites of reduced ATAC-seq signal are luminal-specific. GSEA was carried out on all genes (left) or genes mapped to ATAC-seq peaks that changed at least 2-fold (right) using either the Bas-ECJ or Lum(M)-ECJ genesets. NES and Padj values are indicated below the enrichment plots. **C,** Differential ATAC-seq peaks are enriched for epigenomic features and TF binding. Published ChIP-seq data (Jiang, Wang et al. 2019) for the indicated features in MCF7 cells were plotted for all of the ATAC-seq peaks that were decreased at least 2-fold after CPI-1612 treatment. **D,** Sites of differential ATAC-seq signal are more open in non-basal relative to basal breast tumors. Average ATAC-seq signal across all TCGA samples annotated as either basal or non-basal breast cancer was calculated for each of the differential ATAC-seq peaks described in **Fig 3** and was compared to the difference in the average signal across a set of non-differential ATAC-seq peaks. P-values were calculated by unpaired student’s t-test with Welch’s correction on log-normalized read counts in each peak.

FOXA1 binding sites are known to define lineage-specific enhancers in luminal breast cancer cells (Zhang 2016) and several luminal-specific transcription factor motifs (GATA3, TRPS1, etc.) (Lin 2017) were also significantly enriched in the set of differentially accessibly ATAC-seq sites (Fig 4A). We hypothesized that the enhancers most overtly affected by CPI-1612 were enriched in lineage-specific regulatory elements affecting the expression of luminal gene sets. To test this hypothesis, we used two breast cancer defined gene sets: luminal (Lum(M)-ECJ) to define lineage-specific genes and basal (Bas-ECJ) as a foil for non-luminal genes (Bernardo, Bebek et al. 2013). GSEA of the differentially expressed genes identified from RNA-seq of CPI-1612 treated MCF7 cells revealed significant enrichment of the luminal gene set for down-regulated genes (p=1E-10), while the basal gene set showed no significant enrichment (Fig 4B, left, and Figs S7A and S7B). Performing the same enrichment analysis with genes mapped from the ATAC-seq analysis again demonstrated a significant enrichment (p=1E-3) for the luminal set, though only a subset of the differentially accessible peaks could be mapped to genes (Fig 4B, right).

To check whether the differentially accessible sites were occupied by FOXA1 in MCF7 cells we compared these sites using published ChIP-seq data (Jiang, Wang et al. 2019). Confirming what the HOMER analysis predicted, a majority of the differentially accessible sites were bound by FOXA1 in MCF7 cells (Fig 4C). As shown in Fig S5B, 54% of the differential peaks overlapped with a FOXA1 binding site and these peaks were enriched for FOXA1 binding compared to all open peaks (p-value < 1.0E-8, Fisher’s Exact test). These loci also contained a significant overlap with H3K27ac, H3K4me1, and ER binding sites. Not surprisingly, given that these sites were predominately located in non-promoter enhancers, there was minimal overlap with H3K4me3 signal.

To explore the translatability of these findings to human breast cancer, we next correlated sites with differential accessibility in response to CPI-1612 treatment in MCF7 cells with known regions of open chromatin in breast invasive carcinoma samples from The Cancer Genome Atlas (Corces, Granja et al. 2018). We found that the differentially accessible sites upon CPI-1612 treatment were more likely to be accessible in non-basal relative to basal tumors, while randomly selected ATAC-seq peaks showed no difference based on tumor lineage (Fig 4D). Taken together, these data show that CPI-1612 specifically targets intergenic enhancers defined by FOXA1 binding to promote the downregulation of luminal-specific genes in ER positive breast cancer.

## DISCUSSION

CREBBP/EP300 regulate the growth and signaling of normal and cancerous cells by integrating upstream stimuli to tune transcriptional output. There is substantial evidence and compelling rationale to support the key role of these HAT proteins in the treatment of hormone-dependent breast cancer. In this study we show that CREBBP/EP300 HAT inhibition with the potent and selective inhibitor CPI-1612 can inhibit the growth of breast cancer cell lines *in vitro* and impede tumorigenesis *in vivo* at well tolerated doses with demonstrated target engagement. We further demonstrate that CREBBP/EP300 HAT inhibition acts in an overlapping but nonidentical manner to the standard of care Fulvestrant, supporting its potential development in combination or resistance settings in ER+ breast cancer.

Despite strong evidence linking CREBBP/EP300 to pathological transcriptional programs in cancer, prior to the late 2010s, inhibitors targeting CREBBP/EP300 acetyltransferase activity were limited to cell impermeable substrate analogs (e.g. Lys-CoA), nonselective natural products (e.g. curcumin), or reactive synthetic compounds (e.g. C646)(Dancy and Cole 2015). Work published by AbbVie was the first to conclusively demonstrate the feasibility of CREBBP/EP300 acetyltransferase inhibition with drug-like small molecules, of which A-485 was the first example (Lasko, Jakob et al. 2017). CPI-1612 was identified as a chemically differentiated acetyl-CoA competitive inhibitor with selectivity over other HAT families, superior potency relative to A-485 in both catalytic domain and full-length biochemical assays, and low nanomolar potency in cell-based target engagement assays (Wilson, Patel et al. 2020). ADME properties are also improved relative to A-485, and it should be noted that CPI-1612 is highly brain penetrant and thus suitable for in vivo studies of CNS malignancies.

Phenotypic profiling of CPI-1612 in breast cancer cell lines showed activity across both ER+ and ER-cell lines, with particular sensitivity in several ER+ cell lines. Dependence on the acetyltransferase activity of CREBBP/EP300 in ER+ breast cancer is not surprising given the known physical and functional links between CREBBP/EP300 and ER and has been corroborated in another recently published study (Waddell, Mahmud et al. 2021). However, while Waddell et al. established a link between H3K27 acetylation at enhancers and ER target gene expression, key points were not addressed: comparison of CREBBP/EP300 HAT inhibition with direct ER targeting, the involvement of chromatin accessibility dynamics in transcriptional regulation by CREBBP/EP300, and the features that define the recruitment of CREBBP/EP300 activity to specific loci.

Selective ER degradation by Fulvestrant represents standard of care therapy for ER+ breast cancer. Combination of CPI-1612 with Fulvestrant demonstrated an additive effect on anti-tumor efficacy of a breast cancer xenograft model, arguing for non-overlapping pharmacology of the two mechanisms. In transcriptional profiling experiments, we noted that while both CPI-1612 and Fulvestrant target the ER transcriptional network, their transcriptional effects at the gene level are distinct. At the dose and timepoint explored, direct ER targeting elicited a modest transcriptional response with affected genes defined by the presence of an ERE. While it is not clear from these experiments whether CREBBP/EP300 HAT inhibition amplifies or accelerates the Fulvestrant transcriptional effect or targets a unique set of genes (or a combination of both), we reasoned that the distinct immediate transcriptional impact may be the result of epigenomic remodeling.

A key observation from our epigenomics experiments is the difference between the broad changes in H3K27ac response and the comparatively more subtle changes observed in chromatin accessibility upon CPI-1612 treatment. Similar to published findings, we observed a global reduction in H3K27 acetylation, with downregulated peaks enriched in enhancers relative to transcriptional start sites (Hogg, Motorna et al. 2021, Narita, Ito et al. 2021, Waddell, Mahmud et al. 2021). However, significant changes in chromatin accessibility were much less abundant, but were notably enriched with ER target genes. Other studies of CREBBP/EP300 inhibition show similarly modest changes in chromatin accessibility in both multiple myeloma (Hogg, Motorna et al. 2021) and embryonic stem cells (Narita, Ito et al. 2021), but this phenomenon has not been observed in hormone-dependent cancers. It is not clear from our work whether the changes in chromatin accessibility are a cause or consequence of acute transcriptional silencing of the enhancer-gene pair, though the authors of the above studies propose the latter.

We reason that the loci showing the most pronounced changes in chromatin accessibility upon CPI-1612 treatment represent the sites that are most dependent on CREBBP/EP300 activity. Our observation that these loci tend to be at distal enhancers bound by the pioneer transcription factor FOXA1 provides some insight as to how FOXA1 activates a distinct subset of lineage-defining enhancers (Zhang 2016, Bojcsuk, Nagy et al. 2017, Fu, Pereira et al. 2019). However, while FOXA1 is broadly required for the establishment of open chromatin regions (Hurtado, Holmes et al. 2011, Glont, Chernukhin et al. 2019), our observation that only a small number of sites show acute loss of chromatin accessibility upon CREBBP/EP300 HAT inhibition may imply the existence of additional layers of chromatin regulation to establish or maintain open chromatin such as histone methylation (Jozwik, Chernukhin et al. 2016). Further, it has been shown that ER and FOXA1 show approximately a 50% overlap, arguing for ER binding events that do not directly depend on FOXA1 or CREBBP/EP300 activity and may instead be recruited by features such as “strong” ERE sequences (Hurtado, Holmes et al. 2011, Bojcsuk, Nagy et al. 2017). This differential requirement for FOXA1 and chromatin acetylation/accessibility dynamics for ER chromatin recruitment may contribute to the distinct transcriptional effects of CREBBP/EP300 HAT inhibition relative to direct ER targeting.

From our data it is not known whether FOXA1 directly recruits CREBBP/EP300 activity or whether activity is recruited by additional transcription factors, such as GATA3 or the MegaTrans complex (Cirillo, Lin et al. 2002, Kong, Li et al. 2011, Liu, Merkurjev et al. 2014). Notably, members of the core transcriptional regulatory circuitry in MCF7 such as *GATA3* and *ELF3* (Saint-Andre, Federation et al. 2016) are represented in both the set of differentially accessible chromatin sites and the set of downregulated genes, arguing that CREBBP/EP300 HAT inhibition impacts these interconnected networks on multiple fronts. The role of acetylation dynamics in FOXA1-mediated enhancer activation may be analogous to the recruitment of HDAC activity to leukemia-specific loci by the pioneer transcription factor PU.1 (Frank, Manandhar et al. 2016). Further studies are required to refine the relationship between FOXA1 and CREBBP/EP300 and resolve the temporal relationship among acetylation, chromatin accessibility, and transcription.

This work provides a mechanistic rationale for the exploration of CREBBP/EP300 acetyltransferase inhibition as a therapeutic strategy in ER+ breast cancer. Further studies using the potent and selective inhibitor CPI-1612 will inform clinical development plans and maximize the potential of this approach to address the known limitations of existing therapies that target ER transcription in breast cancer.

## MATERIALS AND METHODS

### Cell lines and compounds

MCF7, T47D, and ZR751 were obtained from ATCC, cultured per supplier’s instructions, and used at early passages for experiments. Cells were routinely screened for mycoplasma contamination. Synthesis of CPI-1612 has been described previously (Wilson, Patel et al. 2020). Fulvestrant (ICI182780) was obtained from Sellekchem for in vitro studies. Fulvestrant (FASLODEX injection/AstraZeneca) was obtained as a dosing solution of 0.25g/5 mL for in vivo studies.

### PRISM cell panel profiling

Pooled screening of barcoded cell lines with a dose titration of CPI-1612 (5 μM, 1.67 μM, 0.6 μM, 0.19 μM, 0.062 μM, 0.021 μM, 0.007 μM, and 0.002 μM) for 5 days was carried out by the Broad Institute PRISM lab according to published protocols (Yu, Mannan et al. 2016). Growth inhibition was assessed using cell barcodes as a proxy for cell number and was expressed as relative growth rate by comparing the cell abundance at the start of the experiment and normalizing to growth rate of the DMSO control. GI_50_ values were calculated using GraphPad PRISM curve fitting to interpolate the concentration at which relative growth rate was 0.5.

### In vitro cell growth assays

For treatments in full serum, cells were plated in 96-well plates and treated in duplicate wells with a dose titration of CPI-1612 for 4 days. For treatments in charcoal-stripped serum, cells were plated in 96-well plates, washed after overnight incubation, and moved to phenol-red free media (Gibco) + 10% charcoal-stripped serum (Gibco) for 2 days. 17-β-estradiol at 100 nM was added along with a dose titration of CPI-1612 for 4 days. Viability was assessed using Cell Titer Glo (Promega), and GraphPad Prism curve fitting was used to fit the data.

### In vivo efficacy studies

All animal studies were carried out at Wuxi AppTec (Shanghai) Co. Inc., in accordance with the Institutional Animal Care and Use Committee (IACUC) of WuXi AppTec following the guidance of the Association for Assessment and Accreditation of Laboratory Animal Care (AAALAC). CPI-1612 was formulated in DMSO/PEG400/H_2_0, v/v/v, 1/3/6. Stock solutions were prepared for dosing at 0.25 and 0.5 mg/kg at a dosing volume of 10 μL/g twice per day by oral gavage. Fulvestrant was delivered at 1 mg/kg at a dosing volume of 20 μL per mouse once per week by subcutaneous injection.

For efficacy studies, female Balb/c mice at 6-8 weeks old were implanted in the left flank with 17β-Estradiol (0.18 mg) pellets (Innovative Research of America Cat. No.: SE-121, pellet size: 3.0 mm). After 4 days, mice were inoculated subcutaneously in the right flank with 1E7 exponentially growing MCF7 cells in 0.2 mL of PBS/Matrigel at a 1:1 ratio. After average tumor volume reached 150 mm^3^, mice were randomized for dosing initiation, with 8 mice per group. Animals were monitored for body weight change or other abnormal effects as stated in the approved protocols, and any animal with deteriorating condition or tumor size greater than 3000 mm^3^ was euthanized. Remaining animals were dosed for 21 days, with tumors measured three times per week in two dimensions using a caliper. Tumor volumes were calculated as V= 0.5a x b^2^, where a is the short diameter and b is the long diameter.

### Plasma pharmacokinetics analysis

At study endpoint, approximately 50 μL blood was collected from 4 mice in each group and placed in EDTA-2K tubes (1.5 ml tube containing 3 μL of 0.5M EDTA-2K). Anticoagulant blood was centrifuged at 2,000 x g at 4 ºC for 15 min. Plasma was stored at −80 ºC before analysis. Plasma samples were analyzed by LC/MS/MS, and concentrations of CPI-1612 and Fulvestrant were determined by comparing to standards.

### PBMC preparation and FACS pharmacodynamics assay

At study endpoint, 300-400 μL of whole blood was collected for PBMC isolation from all mice in each group by Ficoll-Paque media density centrifugation. For FACS staining, 2E5 cells were washed twice with DPBS, centrifuged at 400xg for 5 minutes at room temperature (RT), and the pellet was resuspended by flicking. Live/dead viability dye (Biolegend 423114) was diluted 1:1000 in DPBS, and 100 μL was added to the cell pellet. The plate was incubated for 20 min at RT in the dark. Cells were washed twice with 200 μL FACS staining buffer (Ebiosiences 00-4222-26), centrifuged at 400xg for 5 min at RT, and resuspend in 45 μL staining buffer by flicking. Fc Block (BD Biosciences 553142) was added at 5 μL/well and plate was incubated for 5 min at RT in the dark. Staining buffer (50 μL) was added to each well and the plate was incubated for 30 min at 4°C in the dark. Cells were washed twice with 200 μL staining buffer.

FoxP3 and Transcription Factor Staining Buffer Set (Ebioscience 00-5523-00) was used for intracellular staining. Fixation/permeabilization buffer was diluted to 1x using assay diluent, and 100 μL was added to each well with mixing by pipetting. Plate was incubated for 45 min at 4°C in the dark, 100 μL of 1x permeabilization buffer (1:10 in water) was added, and plate was centrifuged at 450xg for 5 min. Cells were washed once with 200 μL permeabilization buffer, resuspended in 100 μL staining buffer, and stored at 4°C overnight. Cells were blocked by adding 100 μL of 1:1000 rabbit gamma globulin (Jackson Immunoresearch) and incubated for 5 min at RT. Cells were washed with 200 μL permeabilization buffer. PE-conjugated rabbit anti-H3K27ac (Cell Signaling 11562S) was diluted 1:20 in permeabilization buffer, and 100 μL was added to cells and mixed by pipetting. Following a 45 min incubation at 4°C, cells were washed twice with 200 μL permeabilization buffer followed by centrifugation at 450xg for 5 min. Pellet was resuspended in staining buffer and used for FACS analysis. At least 1E4 events (gated on singlet live cells) were acquired for each sample.

### Tumor pharmacodynamics analysis

Tumor samples at study endpoint from same animals as were used for PK analysis were snap frozen in liquid nitrogen and stored at −80°C. Frozen tumor samples were pulverized using a Covaris tissue pulverizer, returned to liquid nitrogen, and stored on dry ice. For analysis of gene expression, samples were resuspended in 1 mL Trizol (Invitrogen) and homogenized with an Omni Tissue Master homogenizer. Homogenized tumors were centrifuged for 10 min at 4°C, and supernatant was transferred to fresh tubes. Total RNA was extracted and precipitated according to Trizol manufacturer’s instructions. First strand cDNA was prepared using SuperScript IV reverse transcriptase (Invitrogen) according to manufacturer’s instructions. Expression of *MYC* in tumors was measured by q-RTPCR with UPL chemistry (Roche, probe #34; F primer 5’tgaattagaatctcgggagtgc3’, R primer 5’ gagtgagaccccatctcagaa3’), and was normalized to the expression of *ACTB* measured by Taqman chemistry (Applied Biosystems #Hs99999903_m1). RT-PCR data were acquired with a LightCycler 480 (Roche).

### Bulk RNA-seq

Cells were treated with CPI-1612 for six hours at 5 or 50 nM with or without 100nM Fulvestrant. RNA was extracted with the RNeasy kit (Qiagen), in accordance with the manufacturer’s protocol. Samples were treated with DNase and polyadenylated (polyA+) RNA was isolated. Sequencing libraries were constructed using the Illumina TruSeq RNA Sample Preparation Kit (v2). The resulting libraries were sequenced on an Illumina HiSeq 4000, with 9 samples multiplexed per lane. 2×150 base pair paired-end reads using the Illumina TruSeq strand specific protocol, for an expected 20 million reads per sample.

### RNA-seq bioinformatics analysis

Isoform expression of Ensembl transcripts (GRCh38 release 99) was calculated with Salmon version 0.11.3 (Patro, Duggal et al. 2017). Gene-level counts were then imported into R using tximport (Soneson, Love et al. 2015). Differential expression analysis was performed with DESeq2 (Love, Huber et al. 2014) to analyze for differences between conditions, and the set of differentially expressed genes were filtered only for the genes with a | fold-change | greater than 1.5 and a Benjamini-Hochberg FDR adjusted P values less than or equal to 0.05.

### ATAC-seq library preparation and sequencing

MCF7 cells were treated with either 50 nM CPI-1612 or DMSO for 6 hours to profile accessible chromatin. Approximately 10,000 cells for each replicate were profiled using the Omni-ATAC-seq protocol as described (Corces, Trevino et al. 2017). Cells were thawed quickly in a 37°C rocking bath and 900 μL of ice-cold PBS supplemented with Roche Complete Mini Protease inhibitor was added immediately. Cells were split into two 1.5-ml Eppendorf DNA lo-bind tubes to serve as technical replicates. Cells were centrifuged at 500xg for 5 min at 4°C, washed once in PBS with protease inhibitor, centrifuged at 500xg for 5 min at 4 °C and supernatant was removed completely using two separate pipetting steps with extreme caution taken to avoid resuspension. The transposition reaction consisted of 20-μL total volume of the following mixture (10 μL 2× TD Buffer, 1 or 0.5 μL TDEnzyme, 0.1 μL of 2% digitonin, 0.2 μL of 10% Tween 20, 0.2 μL of 10% NP40, 6.6 μL of 1× PBS and 2.3 μL of nuclease-free water). Tagmented DNA was purified by QIAGEN MinElute Clean up Kit, PCR amplified and libraries were purified with 1.2 volume of AMPure XP beads. DNA bound to the beads was washed twice in 80% ethanol and eluted in 20 μL of water. Indexed fragments were checked in concentration by qPCR, profiled by Bioanalyzer, equimolarly pooled and sequenced on an Illumina Hi-Seq 4000 with 2 samples multiplexed per lane using 2×150 bp sequencing to a target depth of 60 million reads per sample.

### Bulk ATAC-seq bioinformatics analysis

Paired-end ATAC reads were trimmed to remove Nextera adaptors using Atropos v1.1.28 (Didion, Martin et al. 2017). Trimmed reads were mapped to the human reference hg38 using BWA mem v0.7.17 (Li and Durbin 2009) with default settings. Aligned bam files were filtered with bamtools (Barnett, Garrison et al. 2011) to remove a) reads that are not within 2kb on the same chromosome and b) reads with more than four mismatches to reference. Peaks of open chromatin were identified using Genrich (Gaspar 2020) in ATAC-seq mode (-j) to adjust for Tn5 shift, using its internal PCR duplicate remover (-r), ignoring mitochondrial (-e chrM) and black-listed regions (-E). The bed file of significant peaks was then processed by HINT (Li, Schulz et al. 2019) to identify footprints with Tn5 bias correction (--atac-seq) and HOMER’s (Heinz, Benner et al. 2010) findMotifs.pl script was used to identify enriched motifs in the footprints. using Bins Per Million (bpm) with deepTools (Ramirez, Ryan et al. 2016).

### Single cell ATAC-seq

Following 6 hour treatment with DMSO or 50 nM CPI-1612, single MCF7 cells were prepared in accordance with the 10X Genomics manufacturer’s protocol. scATAC-seq libraries were constructed using the 10X Genomics Chromium Single Cell ATAC-seq library kit. The resulting libraries were sequenced on an Illumina HiSeq 4000, with 2 samples multiplexed per lane. 2×250 base pair paired-end reads using the Illumina TruSeq strand specific protocol, for an expected 150 million reads per sample (targeting 50,000 reads/cell).

### Single cell ATAC-seq bioinformatics analysis

All raw base call (BCL) files generated by Illumina sequencers were converted to FASTQ files using cellranger-atac mkfastq function. Read filtering and alignment, barcode counting, and identification of transposase cut sites were detected using cellranger-atac count function (cellranger-atac_version 1.1.0) using the CellRanger reference package “refdata-cellranger-atac-GRCh38-1.1.0” with “gencode.v28.basic” annotations and Signac v(1.3) (Stuart, Srivastava et al. 2021) while filtering samples for cells with at least 3,000 and less than 20,000 fragments per cell. DMSO and CPI-1612 samples were integrated with Harmony v1.0 (Korsunsky, Millard et al. 2019) and transcription factor activity was calculated with chromVAR v1.12 (Schep, Wu et al. 2017) using the JASPAR2020 motif dataset.

### ChIP-seq

Chromatin was prepared by Diagenode ChIP-seq Profiling service (Diagenode Cat# G02010000) using the iDeal ChIP-seq kit for Histones (Diagenode Cat# C01010059).

Chromatin was sheared using Bioruptor® Pico sonication device (Diagenode Cat# B01060001) combined with the Bioruptor® Water cooler for 6 cycles using a 30’’ [ON] 30’’ [OFF] settings. Shearing was performed in 1.5 ml Bioruptor® Pico Microtubes with Caps (Diagenode Cat# C30010016) with the following cell number: 1 million in 100μL. An aliquot of this chromatin was used to assess the size of the DNA fragments obtained by High Sensitivity NGS Fragment Analysis Kit (DNF-474) on a Fragment Analyzer™ (Agilent).

ChIP was performed using IP-Star® Compact Automated System (Diagenode Cat# B03000002) following the protocol of the aforementioned kit. Chromatin corresponding to 6μg was immunoprecipitated using the following antibodies and amounts: H3K9ac (C15410004 lμg), H3K27ac (C15410196, 1μg). Chromatin corresponding to 1% was set apart as Input. qPCR analyses were made to check ChIP efficiency using KAPA SYBR® FAST (Sigma-Aldrich) on LightCycler® 96 System (Roche) and results were expressed as % recovery = 2^(Ct_input-Ct_sample). Primers used were the following: EIF4a2 prom, GAPDH prom, MyoEx2.

The library preparation has been conducted by Diagenode ChIP-seq/ChIP-qPCR Profiling service (Diagenode Cat# G02010000). Libraries were prepared using IP-Star® Compact Automated System (Diagenode Cat# B03000002) from input and ChIP’d DNA using MicroPlex Library Preparation Kit v2 (12 indices) (Diagenode Cat# C05010013). Optimal library amplification was assessed by qPCR using KAPA SYBR® FAST (Sigma-Aldrich) on LightCycler® 96 System (Roche) and by using High Sensitivity NGS Fragment Analysis Kit (DNF-474) on a Fragment Analyzer™ (Agilent). Libraries were then purified using Agencourt® AMPure® XP (Beckman Coulter) and quantified using Qubit™ dsDNA HS Assay Kit (Thermo Fisher Scientific, Q32854). Finally their fragment size was analyzed by High Sensitivity NGS Fragment Analysis Kit (DNF-474) on a Fragment Analyzer™ (Agilent). 2×150 bp paired-end reads were sequenced on an Illumina HiSeq 2500 with a target depth of 60 million reads per sample.

### ChIP-seq bioinformatics analysis

Paired-end ChIP reads were trimmed to remove Illumina adaptors using Atropos v1.1.28 (Didion, Martin et al. 2017). Trimmed reads were mapped to the human reference hg38 using BWA mem v0.7.17 (Li and Durbin 2009) with default settings. Aligned bam files were filtered with bamtools (Barnett, Garrison et al. 2011) to remove a) reads that are not within 2kb on the same chromosome and b) reads with more than four mismatches to reference. PCR duplicate reads were removed using Picard (http://broadinstitute.github.io/picard/). Peaks of H3K27ac and H3K9ac were identified using sample-matched input as a control with SICER2 (Xu, Grullon et al. 2014) with the gap size (-g) equal to 600 bp. Genomic bigwig tracks were normalized using Bins Per Million (bpm) with deepTools (Ramirez, Ryan et al. 2016).

## ACKNOWLEDGEMENTS

The authors would like to thank colleagues at Constellation for thoughtful discussions and review of the manuscript.

## DATA AVAILIBILITY

Datasets generated during this study are deposited at the Gene Expression Omnibus (GEO): GSE190163.

## AUTHOR CONTRIBUTIONS

A.B-R. and S.P-C. designed and carried out experiments and analyzed data. D.N.M. and M.J.S. analyzed next generation sequencing data. E.T. and A.H. carried out experiments. J.E.W. and R.J.S. 3^rd^ contributed to experimental design. A.R.C. analyzed data. A.B-R., S.P-C., D.N.M, and A.R.C. wrote the manuscript.

