## Supplementary Figure Legends for "CREBBP/EP300 acetyltransferase inhibition disrupts FOXA1-bound enhancers to inhibit the proliferation of ER+ breast cancer cells"

**Figure S1 (related to Figure 1). A,** Pooled, barcoded cell lines were treated with a dose titration of CPI-1612 for 4 days, and growth inhibition was calculated using depletion of cell line barcodes relative to initial representation. The concentration at which growth was inhibited to 50% of the untreated cells (GI_50_) was calculated. Red, ER+ cell lines; black: ER- cell lines. **B,** As in **Figure 1A**, except cells were treated in media containing charcoal-stripped serum and added estradiol. Error bars represent SD of 2 replicates. **C,** Change in body weight relative to dosing initiation during treatment with CPI-1612. Error bars represent the SEM at each time point. **D,** Plasma concentration of CPI-1612 at study endpoint. Data are expressed as mean and SEM across 4 mice, and p-value was calculated using student’s t-test. **E,** Change in body weight relative to dosing initiation during treatment with CPI-1612 or Fulvestrant. Error bars represent the SEM at each time point. **F,** Plasma concentration of CPI-1612 at study termination for single agent or combination treatment. Data are expressed as mean and SEM across 4 mice, and p-value was calculated using student’s t-test. *ns:* p>0.5. **G,** Plasma concentration of Fulvestrant at study termination for single agent or combination treatment. Data are expressed as mean and SEM across 4 mice, and p-value was calculated using student’s t-test. *ns:* p>0.05.

**Figure S2 (related to Figure 2). A,** Venn diagram of genes down- or upregulated by CPI-1612 or Fulvestrant treatment as in **Figure 2**. Numbers indicate genes down- or upregulated at least 1.5-fold with an adjusted p-value <0.05 in DESeq2 comparisons to DMSO-treated cells. **B,** Summary of GSEA against Hallmark genesets for MCF7, T47D, and ZR751 cells treated with the indicated compounds as described in **Figure 2**. **C,** Enrichment plots for GSEA of RNA-seq data for the HALLMARK_ESTROGEN_RESPONSE_EARLY geneset in MCF7, T47D, or ZR751 cells treated with CPI-1612, Fulvestrant, or CPI-1612 + Fulvestrant as described in **Figure 2**.

**Figure S3 (related to Figure 2). A,** Example of differential gene regulation by CPI-1612 and Fulvestrant in T47D and ZR751 cells as described in Figure 2D. **B,** Fraction of differentially expressed genes with an annotated Estrogen Response Element (ERE) as previously defined (Lin, Vega et al.) with estrogen-responsive genes in MCF7 cells with an adjacent ChIP-identified ER binding site.

**Figure S4 (related to Figure 3).** Comparison of changes in chromatin accessibility and H3K27ac after treatment with CPI-1612. **A**, Venn diagram showing the overlap of all ATAC-seq peaks H3K27ac peaks. **B,** Table summarizing total ATAC-seq and H3K27ac peaks, and peaks with a 2-fold change upon CPI-1612 treatment. **C,** Heatmap of differential ATAC-seq peaks showing that peaks that show the largest change in ATAC-seq signal (k-means cluster #1) are most likely to show a reduction in H3K27ac signal. **D,** Fraction of differential ATAC-seq peaks that are also differential H3K27ac peaks (blue bar), and fraction of differential H3K27ac peaks that are also differential ATAC-seq peaks (orange bar).

**Figure S5 (related to Figure 4). A,** HOMER motif search of differential H3K27ac peaks upon CPI-1612 treatment. Binding sites for FOXA1 and luminal specific TFs are not enriched in differential H3K27ac peaks relative to all H3K27ac peaks. **B,** Overlap of transcription factor (TF) binding with all identified ATAC-seq peaks or those peaks that showed at least a 2-fold reduction in signal after CPI-1612 treatment. ER and FOXA1 data are as described in **Figure 4C**.

**Figure S6 (related to Figure 4). Single cell ATAC-seq after CPI-1612 treatment in MCF7 cells. A,** UMAP dimensionality reduction plot of scATAC-seq data colored by treatment for both unintegrated cells and data integrated using Harmony (Korsunsky, Millard et al.). **B,** Waterfall plot of peaks identified from scATAC-seq data ranked by log_2_ (fold-change) for CPI-1612 relative to DMSO. Blue, peaks reduced by at least 2-fold. **C,** UMAP plot as in **A,** colored by predicted FOXA1 activity based on ChromVAR analysis. **D,** Quantification of predicted FOXA1 activity for DMSO and CPI-1612 treated cells. Boxplots depict median and range of FOXA1 (MA0148.4 motif) activity; p-value was calculated with the Mann-Whitney U test.

**Figure S7 (related to Figure 4). A,** Differential expression of genes in the Bas-ECJ or Lum(M)-ECJ gene sets upon treatment with CPI-1612. **B,** Volcano plot of gene expression changes as described in **B**.
