## Supplementary figures and images for "CREBBP/EP300 acetyltransferase inhibition disrupts FOXA1-bound enhancers to inhibit the proliferation of ER+ breast cancer cells"

### Figure S1

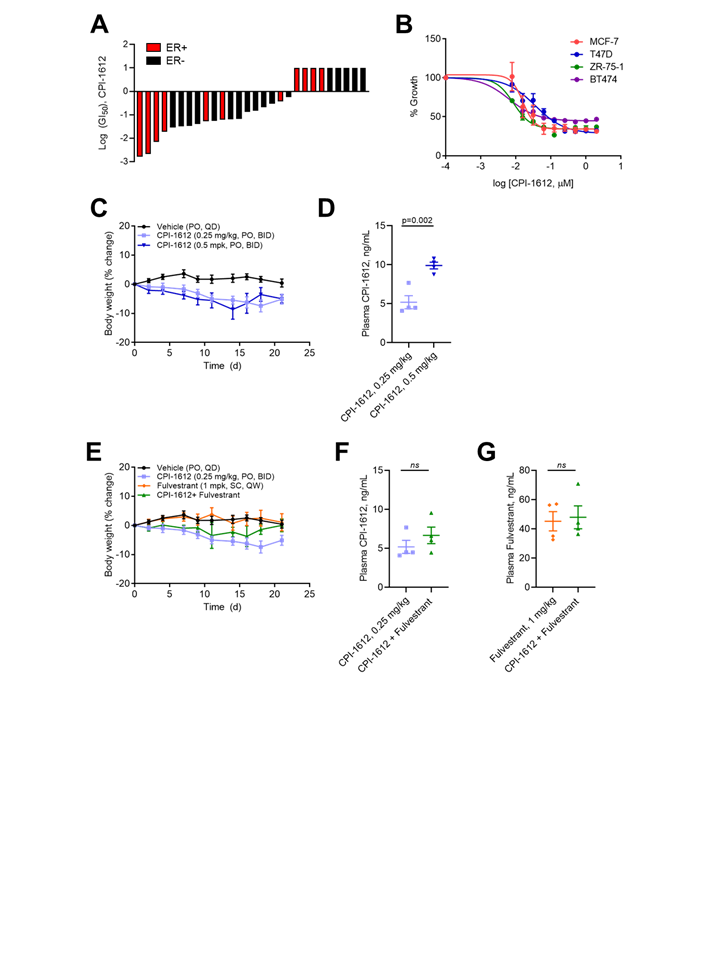

### Figure S2

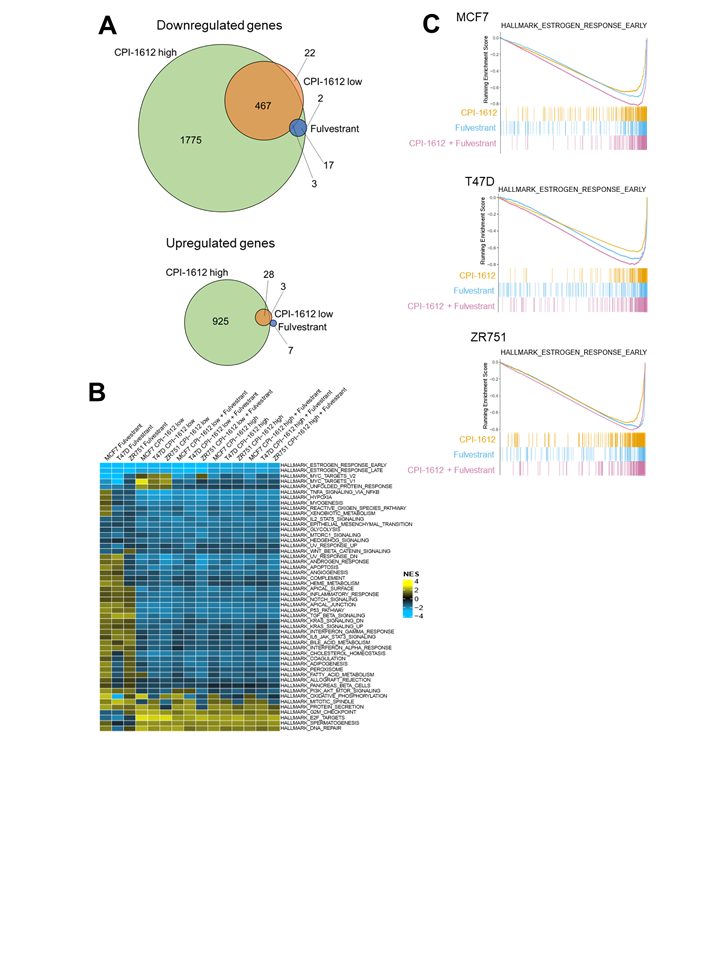

### Figure S3

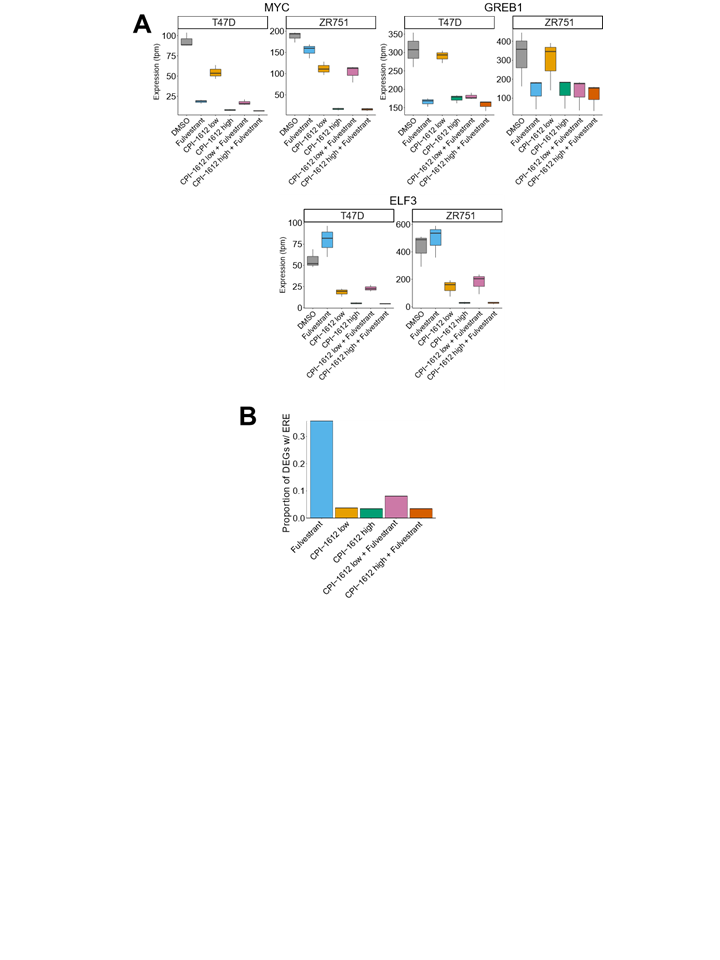

### Figure S4

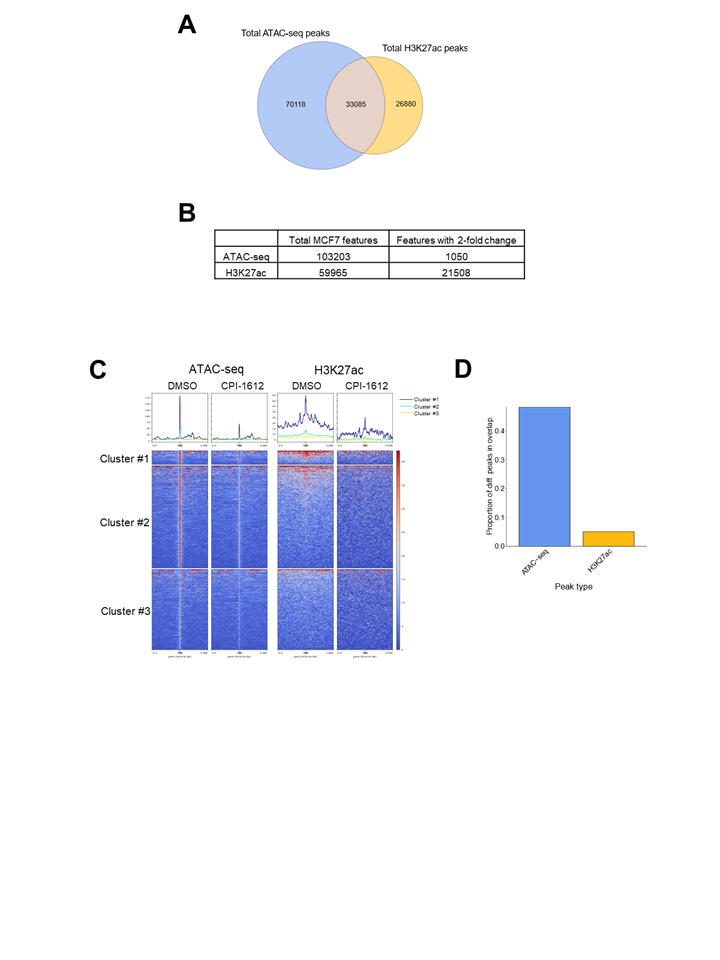

### Figure S5

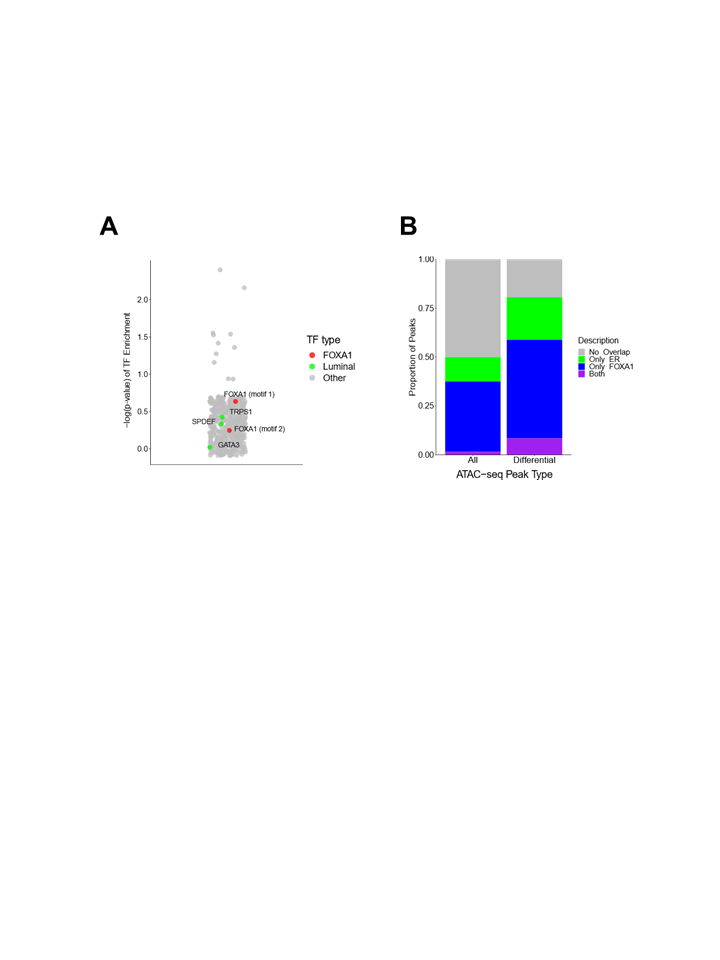

### Figure S6

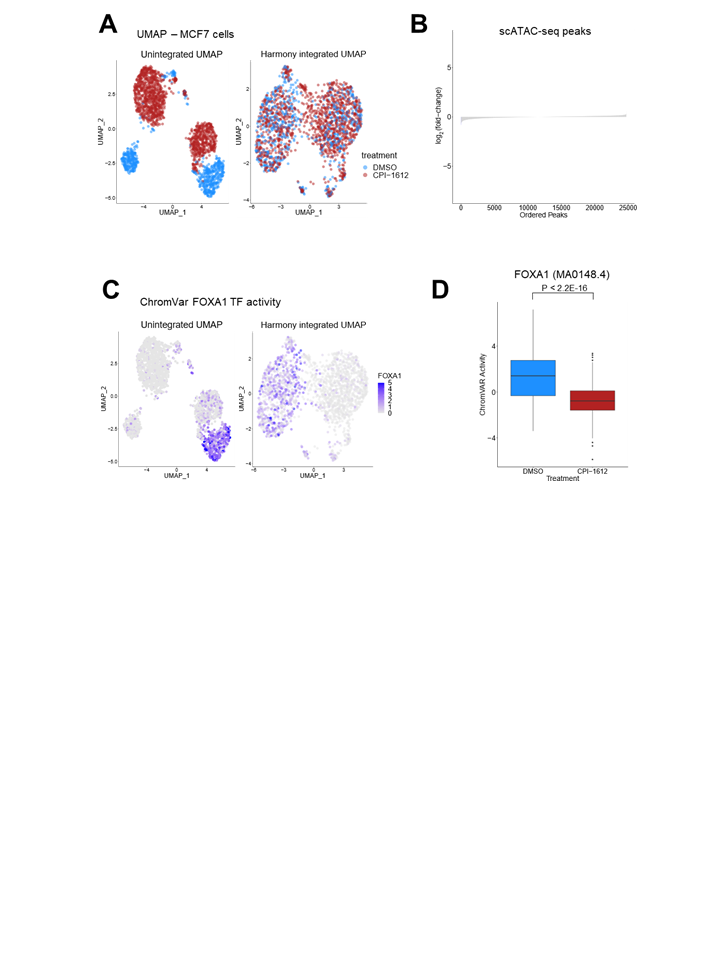

### Figure S7

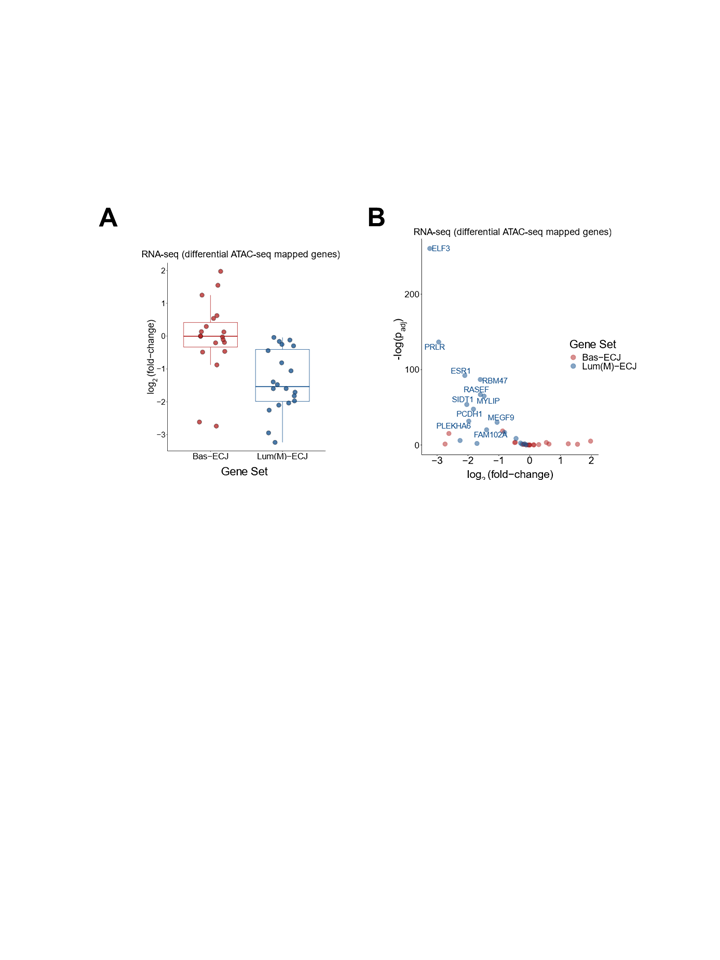
